# Bedscales: A non-contact adherence-independent multi-person sensor for longitudinal physiologic monitoring in the home bed

**DOI:** 10.1101/2020.03.31.018523

**Authors:** Nicholas Harrington, Zhe Wei, Brandon Hernandez-Pacheco, Quan Bui, Pamela N. DeYoung, Bayan Duwaik, Akshay S. Desai, Deepak L. Bhatt, Robert L. Owens, Todd Coleman, Kevin R. King

## Abstract

Home health monitoring technologies promise to improve care and reduce costs, yet they are limited by the need for adherence to self-monitoring, usage of an app, or application of a wearable. While implantable sensors overcome the adherence barrier, they are expensive and require invasive procedures. Here, we describe a non-invasive, non-contact, adherence-independent sensor, that when placed beneath the legs of a patient’s home bed, longitudinally monitors total body weight, detailed respiratory signals, and ballistocardiograms for months, without requiring any active patient participation. Accompanying algorithms demix weight and respiratory signals when the bed is shared by a partner or a pet. Validation studies during overnight clinical sleep studies exhibit quantitative equivalence to commercial sensors and allow discrimination of obstructive and central sleep apneas. In-home studies discriminate atrial fibrillation from normal sinus rhythm. To demonstrate real-world feasibility, we performed 3 months of continuous in-home monitoring in a patient with heart failure as he awaited and recovered from coronary artery bypass surgery. By overcoming the adherence barrier, Bedscales has the potential to create a multidimensional picture of chronic disease, learn signatures of impending hospitalization, and enable optimization of care in the home.

**Disclosures:** Drs. King and Coleman and Nicholas Harrington are inventors on a patent application describing the Bedscales technology. Dr. Kevin R. King discloses consulting relationships with Bristol Myers Squibb and Astrazeneca, Medimmune and is founder of Nightingale Labs. Dr. Deepak L. Bhatt discloses the following relationships - Advisory Board: Cardax, Cereno Scientific, Elsevier Practice Update Cardiology, Medscape Cardiology, PhaseBio, Regado Biosciences; Board of Directors: Boston VA Research Institute, Society of Cardiovascular Patient Care, TobeSoft; Chair: American Heart Association Quality Oversight Committee; Data Monitoring Committees: Baim Institute for Clinical Research (formerly Harvard Clinical Research Institute, for the PORTICO trial, funded by St. Jude Medical, now Abbott), Cleveland Clinic (including for the ExCEED trial, funded by Edwards), Duke Clinical Research Institute, Mayo Clinic, Mount Sinai School of Medicine (for the ENVISAGE trial, funded by Daiichi Sankyo), Population Health Research Institute; Honoraria: American College of Cardiology (Senior Associate Editor, Clinical Trials and News, ACC.org; Vice-Chair, ACC Accreditation Committee), Baim Institute for Clinical Research (formerly Harvard Clinical Research Institute; RE-DUAL PCI clinical trial steering committee funded by Boehringer Ingelheim; AEGIS-II executive committee funded by CSL Behring), Belvoir Publications (Editor in Chief, Harvard Heart Letter), Duke Clinical Research Institute (clinical trial steering committees, including for the PRONOUNCE trial, funded by Ferring Pharmaceuticals), HMP Global (Editor in Chief, Journal of Invasive Cardiology), Journal of the American College of Cardiology (Guest Editor; Associate Editor), Medtelligence/ReachMD (CME steering committees), Population Health Research Institute (for the COMPASS operations committee, publications committee, steering committee, and USA national co-leader, funded by Bayer), Slack Publications (Chief Medical Editor, Cardiology Today’s Intervention), Society of Cardiovascular Patient Care (Secretary/Treasurer), WebMD (CME steering committees); Other: Clinical Cardiology (Deputy Editor), NCDR-ACTION Registry Steering Committee (Chair), VA CART Research and Publications Committee (Chair); Research Funding: Abbott, Afimmune, Amarin, Amgen, AstraZeneca, Bayer, Boehringer Ingelheim, Bristol-Myers Squibb, Chiesi, CSL Behring, Eisai, Ethicon, Ferring Pharmaceuticals, Forest Laboratories, Fractyl, Idorsia, Ironwood, Ischemix, Lilly, Medtronic, PhaseBio, Pfizer, PLx Pharma, Regeneron, Roche, Sanofi Aventis, Synaptic, The Medicines Company; Royalties: Elsevier (Editor, Cardiovascular Intervention: A Companion to Braunwald’s Heart Disease); Site Co-Investigator: Biotronik, Boston Scientific, CSI, St. Jude Medical (now Abbott), Svelte; Trustee: American College of Cardiology; Unfunded Research: FlowCo, Merck, Novo Nordisk, Takeda. Dr. Akshay S. Desai discloses the following relationships – Research grants to Brigham and Women’s Hospital to support clinical trial activities from Alnylam, AstraZeneca, and Novartis; Consulting fees from Abbott, Alnylam, AstraZeneca, Biofourmis, Boehringer-Ingelheim, Boston Scientific, Merck, Novartis, Relypsa, Regeneron. Dr. Owens reports consulting fees from Novartis, and research grants to UCSD from Snoozeal, Nitto Denko, and Masimo.

## INTRODUCTION

The human body is in continuous motion. From early in development until the last day of life, our bodies exert a wide range of static and dynamic mechanical forces on the environment. Under most conditions, large musculoskeletal forces obscure lower amplitude physiologic forces resulting from respirations or the ballistic motion of the beating heart. However, when a person is asleep, musculoskeletal motion quiets for extended periods of time, allowing longitudinal monitoring of dynamic physiology for many hours every day within the comfort of a person’s home bed. This is an ideal setting for developing new chronic disease monitoring and management tools.

Cardiopulmonary diseases are among the most challenging chronic conditions to manage. Heart failure (HF), for example, affects 6 million US patients and costs more than $30B per year largely due to the high burden of inpatient care required for management of recurrent exacerbations (*1*). Most hospitalizations for HF are related to worsening congestion that drives symptoms of dyspnea, fatigue, and edema that progressively limit physical activity. It is believed that many of these hospitalizations could be prevented with better strategies for early identification of worsening fluid retention to direct early intervention with pharmacological therapy (*2, 3*). A technology capable of monitoring and learning signatures of early exacerbations and impending hospitalizations would represent a major advance.

Current strategies for remote heart failure management focus on telemonitoring strategies that emphasize regular surveillance of daily weights and changes in vital signs and respiratory symptoms (*4*). These strategies frequently rely on a high degree of patient engagement and self-monitoring, and most studies have accordingly shown little incremental impact on rates of hospitalization and death in HF patients when compared with usual clinic-based care (*5-9*). The need for adherence to self-application of wearables and patient-initiated utilization of apps also limits their generalized utility as home health monitors (*10, 11*). Implantable hemodynamic monitors, such as the CardioMEMS pulmonary artery pressure sensor, have been developed to overcome adherence barriers and provide access to more detailed physiologic data to guide HF management; (*12*), Although use of this technology to augment routine HF care was associated with lower rates of HF hospitalization in the CHAMPION clinical trial, this approach is challenging to scale for population health management given the need for an invasive procedure to implant the sensor and high associated costs, and still requires adherence to a daily schedule of data transmission by the patient(*13, 14*). Newer remote monitoring approaches overcome adherence barriers by leveraging software modifications to existing cardiac rhythm management devices to provide multiparameter monitoring of physiologic signals including heart rate, heart sounds, respiration, physical activity, and intrathoracic impedance that can be integrated to anticipate worsening HF events. While this approach is being tested in clinical trials as a strategy for routine HF management, it is not likely to be helpful for the large proportion of HF patients (particularly those with preserved EF) who do not require implantable pacemakers and defibrillators (*15*). Given limitations of existing remote monitoring technologies for HF, we sought to develop a non-invasive, multiparameter measurement technology that passively collects data without the requirement for daily patient engagement with technology in the home.

Non-contact sensors have been developed using under-the-mattress piezoelectric-, strain gauge-, or radio frequency-based sensing, and are highly sensitive to dynamic respiratory and ballistocardiographic signals (*16, 17*). However, they are sensitive to subject-sensor proximity and orientation and are unable to reliably differentiate signals from individuals who share a bed with a partner or pet (*18-20*). Moreover, because these sensors do not span the entire bed, they are also unable to measure total body weight, which is still the most pervasively-used objective biomarker of worsening congestion that can be quantified in the home.

We developed an under-the-bed mechanical sensing platform (“Bedscales”) that achieves adherence-independent noncontact longitudinal physiological monitoring of total body weight, respirations, and ballistocardiograms (BCG) and tested it in 31 patients with a variety of cardiopulmonary diseases (Table 1). Signals are passively acquired during sleep, without the need for patient participation, even if the bed is shared by a partner or pet. We provide validation of the technology by comparing Bedscales data to ground truth data from commercial bathroom scales and overnight sleep study sensors. We also demonstrate the ability to study pathology by quantitatively characterizing central and obstructive sleep apneas, and by discriminating atrial fibrillation from normal sinus rhythm. Finally, we establish real-world feasibility by performing 3-month, in-home, continuous monitoring of a patient with heart failure before and after coronary artery bypass surgery. By eliminating the need for patient participation, we believe our data provide initial support for use of non-contact home health sensors to improve care of patients with difficult-to-manage chronic cardiopulmonary diseases such as heart failure.

## RESULTS

### Design of a non-contact adherence-independent sensor for chronic disease monitoring

Patients spend more than 90% of their lives at home and greater than 1/3 of their lives in bed. To leverage this optimal diagnostic setting, we designed sensors that could be placed beneath each leg of a conventional home bed, recliner, or couch to perform longitudinal monitoring of total body weight, respirations, and ballistocardiograms without requiring patient participation for self-sensing, data-transmission, or application of a wearable (**Fig. 1A**). Each non-contact low-profile sensor is comprised of force-sensing strain gauges configured in a wheatstone bridge circuit. The signal is amplified and digitized by a 24-bit analog-to-digital integrated circuit with accompanying discrete circuit elements arranged on a compact custom circuit board that snap fits into the custom plastic injection-molded housing (**Fig. 1B**). The housing design includes a planar plastic spring mechanism, which focuses the entire load through the sensing elements and minimizes shunting of force via the surrounding plastic. The assembled device is outfitted with a rubber top and feet to prevent lateral sliding. For in-home applications, a rigid circular bottom plate is added to provide a reliable surface for the sensor feet to contact (**Fig. 1B**). This plate allows the sensors to be used in carpeted bedrooms or outfitted with furniture pads for use on uncarpeted floors. The devices are scalably manufactured using injection molding and are thus suitable for appropriately powered clinical studies (**fig. S1**).

**Figure 1.**
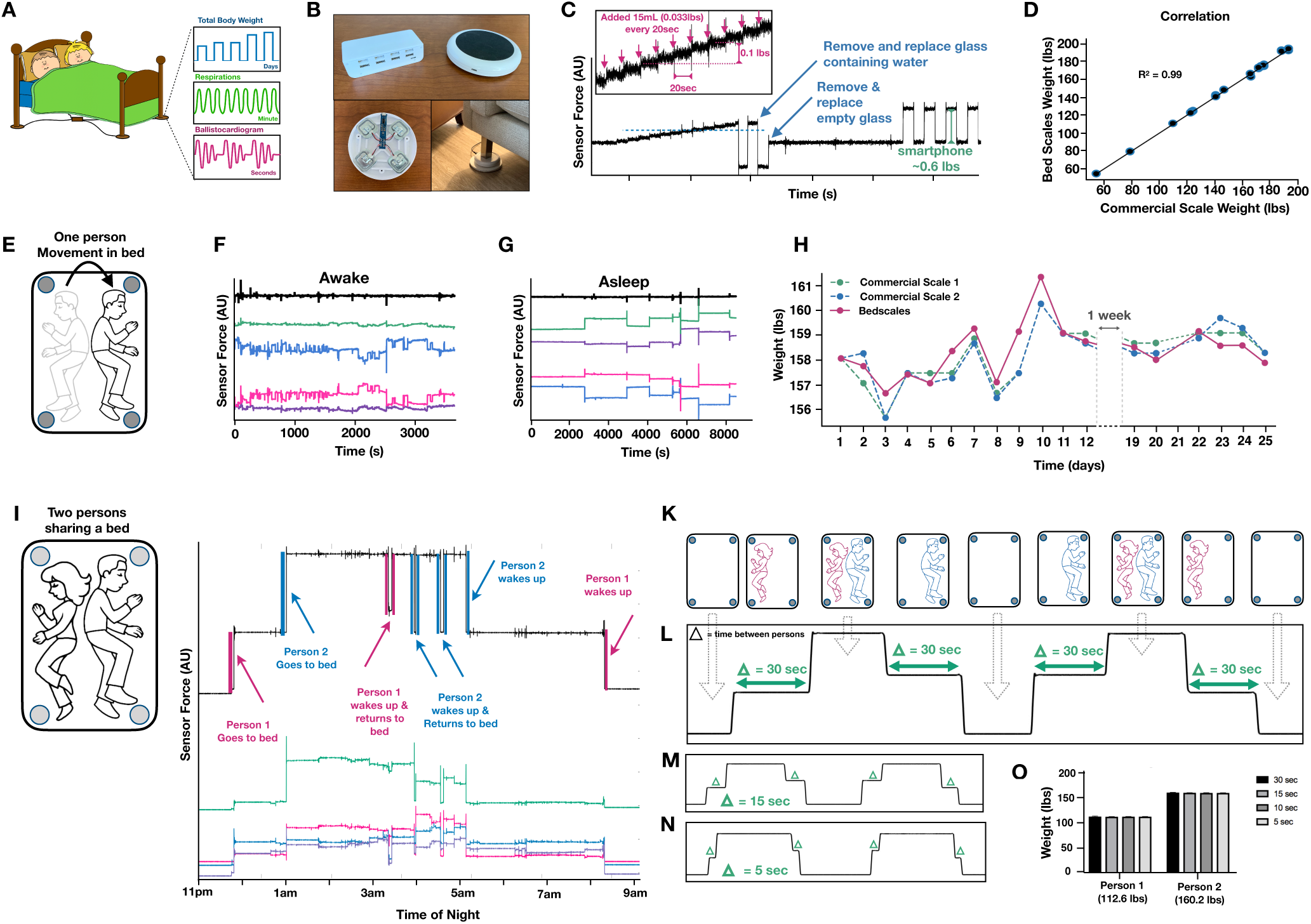
Non-contact adherence-independent multi-person longitudinal weight monitoring using Bedscales. (**a**) Cartoon of the Bedscale concept in which low profile mechanical sensors placed beneath the legs of a conventional home bed measure total body weight, respirations, and ballistocardiogram even if the bed is shared by a partner or pet. (**b**) Pictures of the sensor hardware including communication box (upper left), sensor plastic housing with black rubber tread top (upper right), strain gauges and custom circuit (lower left) and sensor on circular flat plate beneath a furniture leg (lower right). (**c**) Progressive addition of 15mL of water (0.033 lbs) every 20 seconds, followed by removal and replacement of the full and empty glass, followed by addition and removal of a smartphone (0.6 lbs). (**d**) Calibrated Bedscale weights compared to a commercial bathroom scale measuring 19 volunteers getting onto and off of a couch. (**e**) Cartoon illustrating a sleeping subject redistributing total body weight during episodic movements in bed. (**f**) An awake subject using a laptop in bed with frequent redistributions of load (colored signals are individual sensors) yet constant total body weight (black is the sum of sensors). (**g**) A sleeping subject with episodic movements separated by long periods of lying still. (**h**) Comparison of daily Bedscale weight measurements compared to two commercial bathroom scales. (**i**) Cartoon of partners sharing a bed. (**j**) Overnight measurements of two partners sharing a bed. Colored signals indicate individual sensor tracings. Black indicates the sum of sensors. Sudden weight changes due to each person getting into and out of the bed are color coded (blue and pink) and annotated. (**k**) Illustration of synchronicity protocol for measuring total body weights at progressively shorter time intervals. (Person = P) P1 On, P2 On, P1 Off, P2 Off, then repeated exchanging P1 and P2. (**l**) Corresponding total weight signal measured by the Bedscales with a time interval of 30 seconds, (**m**) 15 seconds, and (**n**) 5 sec. (**o**) Estimates of decoupled weights of the two individuals sharing a bed for each synchronicity time interval and corresponding weight measured by a commercial floor scale.

Digitized and amplified data from the individual force sensors is transmitted via micro USB to a central communications module using the open-source Raspberry Pi architecture. We selected a hardwired solution over wireless or Bluetooth so that patients would not have to re-establish communication if Bluetooth pairing is lost. After local signal conditioning, the data is communicated to the HIPAA-affiliate Amazon Web Services via WiFi, or if a local internet connection is unavailable, via a HIPAA-affiliate 3G cellular data transmission device. The measurement system is self-sufficient and does not require an accompanying smartphone or laptop computer. Once stored in the cloud, data can be synchronously or asynchronously processed to create custom analytics, visualizations, and dashboards for permission-dependent sharing with patients, healthcare providers, or family and friends. In the current instantiation, the use of wall power eliminates constraints on the duration of data collection and avoids the need for battery changes. In this way, the system assumes nothing about a patient’s technical literacy or “connectedness” and only requires that a patient have electricity in the home, making it suitable to address management challenges in patients who are socioeconomically disadvantaged, geographically separated from providers, or cognitively impaired.

### Passive in-bed total body weight monitoring

In contrast to bathroom scales, which measure standing weights, Bedscales measure the distributed weight of the bed and its contents such that the total load is proportional to the sum of the individual bed leg measurements. Hospital beds use a similar strategy but do so at a single time point, leaving them vulnerable to errors when objects other than the patient are added between the time of zeroing and measuring weight. We designed Bedscales to perform continuous monitoring, which allows each load to be separately weighed at the time it is added. When an object of constant weight is moved to different locations on the bed, its load redistribution changes but the total remains constant. To calibrate the system at the time of installation, we move an object of known weight (25 lbs) to N+1 locations (where N is the number of bed legs) while making continuous measurements and then assume a linear model to solve for the calibration factors (**fig. S2**). Alternatively, calibration can be performed using a person’s known weight at one time point combined with redistributions of weight during sleep. To explore the lower limits of sensitivity and resolution, we added 15mL aliquots of water (0.033 lbs) every 20 seconds for a total of 4 minutes (180mL, 0.396 lbs) followed by removal and replacement of the full and empty glass, and removal and placement of a smartphone (∼0.6lbs), which are representative of small objects commonly placed on a bed (**Fig. 1C**). This revealed the rms noise to be ∼0.02 lbs (**Fig. 1C**). For comparison, the limit of resolution of conventional bathroom scales is typically 0.2 lbs. When we compared total body weight measurements of healthy volunteers using the Bedscales and a commercial floor scale, we observed strong correlation (R^2^ = 0.99, n=19) (**Fig. 1D**). To validate the measurement of human weight changes that might be experienced during progressive volume overload, we compared the bed sensor to a commercial bathroom scale while a healthy volunteer added and removed increasingly heavy ankle weights before getting into and out of the bed and observed a clear linear relationship (R^2^ = 0.99) (**fig. S3**).

Individuals change position nearly continuously while awake but only episodically during sleep (**Fig. 1E**). **Fig. 1F** illustrates a ∼60-minute segment of data collected while a subject is in bed but awake using a laptop computer. Although the load redistributions are frequent, the total measured weight remains constant. **Fig. 1G** illustrates a ∼3 hour recording during sleep when load redistributions are considerably less frequent. **Fig. 1H** shows a comparison of several weeks of daily weights measured by the Bedscales compared to two commercial floor scales (each with a reported accuracy of ∼0.2 lbs). Errors between the Bedscale and individual commercial scales were comparable to the error between the commercial scales.

### Weight monitoring of multiple individuals sharing a bed

Individuals often share the bed with a partner or pet (**Fig. 1I**); however, they will rarely get into bed at precisely the same time. We reasoned that weights could be separately inferred based on the small delay in the timing of their arrival into bed (**Fig. 1J**). To determine the minimum interval that would allow discrimination of two-person weights, we performed simultaneity tests in which two persons entered and exited the bed at successively decreasing time intervals (**Fig. 1K-O**). Even when the interval was reduced from 30 seconds to 5 seconds, the two individuals were easily discriminated and reproducibly weighed (**Fig. 1O**). Taken together, these data indicate that, compared with a conventional scale, Bedscales can perform high-resolution total body weight measurements in a patient’s home bed, even if the bed is shared with a partner.

### Respiratory monitoring using non-contact in-bed sensors

When a patient is asleep in bed, episodic musculoskeletal movements are separated by comparatively long movement-free intervals during which low variance physiological signals such as respirations and cardiac contractions can be measured with high fidelity. We reasoned that chest wall movements during respirations would generate a detectable shift in the distribution of weight between the sensors beneath each leg. Indeed, we found that frequency-dependent filtering with cutoffs at 0.167 Hz and 1.5 Hz revealed an oscillatory signal characteristic of respirations, consistent with the redistribution of load that accompanies chest wall movement during inspiration and expiration (**Fig. 2A**). We used principle component analysis within a sliding window to calculate eigenvalues that could be multiplied by individual sensor measurements and algebraically summed to create a single optimal respiratory signal with algorithmically-detectable peaks. The resulting signal exhibited brisk linear upstrokes consistent with inspiration, followed by longer exponential decays consistent with expiration. It also allowed quantification of inspiration-expiration I:E ratios, which are prolonged in obstructive lung diseases such as asthma or COPD (**Fig. 2B**) (*21-23*). To validate the measurements, we compared evolving Bedscale respiratory rates to simultaneously recorded commercial respirometry belt measurements (**Fig. 2C**) and observed close correlation with an R^2^ of 0.9992 (**Fig. 2D, fig. S4-5**) and Bland-Altman plots with a standard deviation of ∼0.04 bpm (**Fig. 2E, fig. S4-5**).

**Figure 2.**
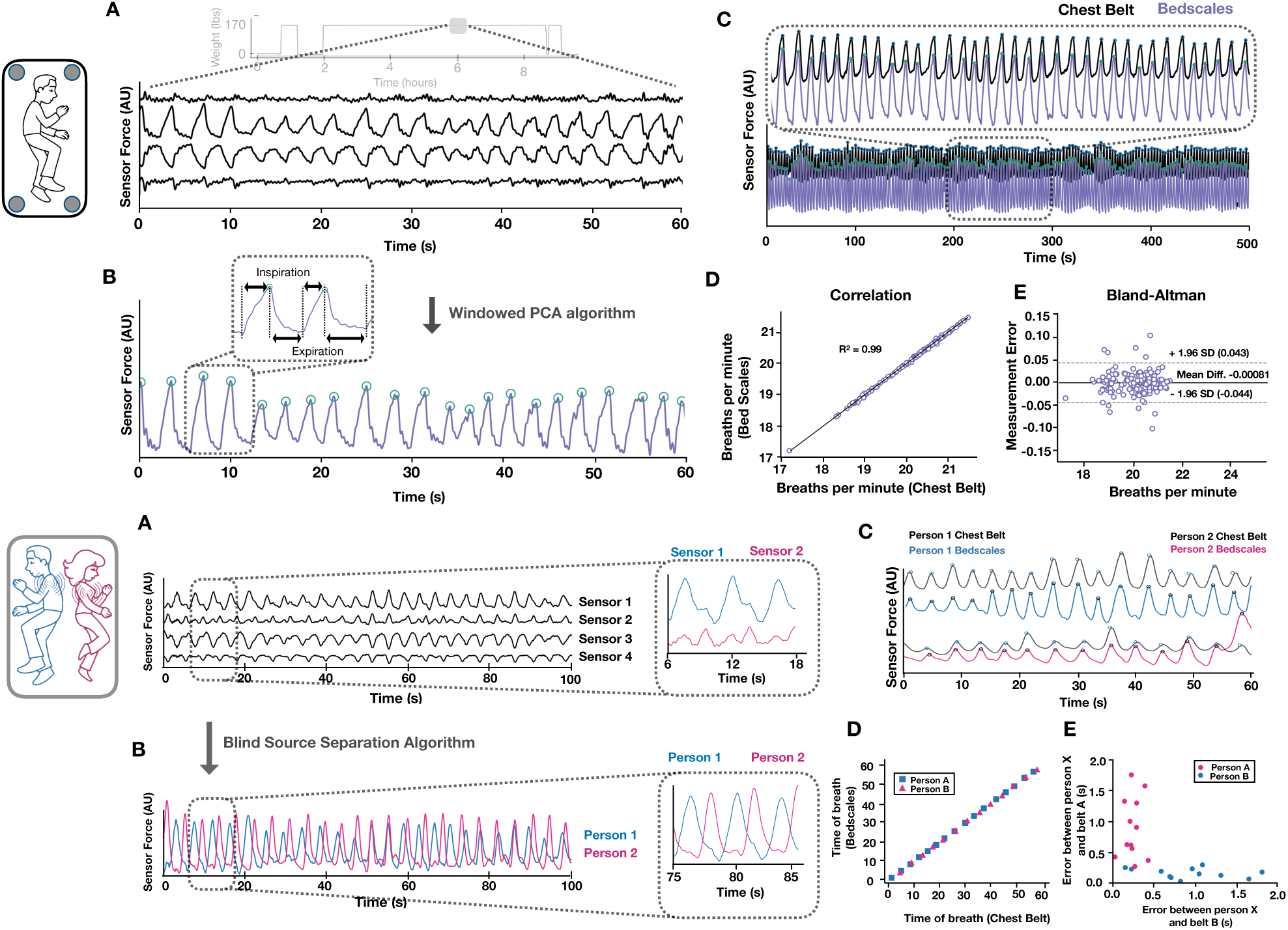
Respiratory monitoring of multiple people in bed using non-contact adherence-independent Bedscales. (**a**) Raw respiratory signals from 4 scales in the middle of an overnight recording (light grey is the entire overnight weight signal). (**b**) Composite signal (purple) derived from linear combination of scales weighted by PCA-based eigenvalues with peak-finding annotation (green). Inset shows short inspiratory phase with rapid linear increase during inspiration followed by longer exponential decay during passive expiration. (**c**) Comparison of commercial respiratory chest belt (black, top signal) compared to the Bedscale respiratory signal (purple, bottom signal) across short (top) and long (bottom) time scales. (**d**) Correlation plot comparing chest belt and Bedscale respiratory rates. (**e**) Bland-Altman plot comparing chest belt and Bedscale respirations. (**f**) Cartoon illustrating how two persons sharing a bed are modeled as two respiratory point sources and raw signals from the 4 legs of the bed beneath two sleeping individuals. Inset shows 2 sensors each predominantly measuring one person with contaminating signal from the second person. (**g**) Demixed signals for person 1 (blue) and person 2 (pink) derived from the raw signals in F. (**h**) Validation experiment comparing demixed Bedscale signals (blue and pink) with the corresponding ground truth chest belt signals (black). (**i**) Correlation of chest belts and the corresponding demixed signal for person 1 and person 2. (**j**) Error between each demixed Bedscale signal and each ground truth respiratory belt signal.

### Demixing respiratory signals of multiple individuals sharing a bed

Existing bed physiology sensors are unable to separate signals from individuals who share the bed or rely on strong assumptions about sidedness, location, and proximity to the sensor. We sought a fully generalizable respiratory demixing solution. We noticed that when an individual person sleeps on the bed, the magnitude of the measured respiratory signal differs between each bed leg but is consistent across time between episodic movements and is position-dependent (**fig. S6**). Therefore, we modeled each person as a respiratory point source (**Fig. 2F**). This allowed the respiratory signals of two individuals sharing the bed to be demixed using source separation mathematics (*24*). We used a hidden Markov model and interpreted the mechanical respiratory sources as latent signals that evolve in a stochastically continuous manner and are mixed through a linear operation with additive sensor noise to give rise to the signals at the individual detectors. Interpreting the linear operation as unknown, we used the expectation-maximization algorithm to find the maximum-likelihood estimate (*25*). Given this estimate, we used the Kalman smoothing algorithm to extract the mechanical respiratory patterns of the two sources. One can see that when two individuals’ sleep in bed at the same time, signals have distinct respiratory patterns that go in and out of phase (**Fig. 2G**). To validate the demixing strategy, we measured Bedscale signals of two individuals sharing the bed while simultaneously recording ground-truth respiratory signals using commercial chest belts. The demixed Bedscale signals strongly correlated with those of the corresponding respiratory belt (**Fig. 2H-J**), showed minimal error when compared to the corresponding person’s chest belt but a large error when compared to the opposite person’s belt data. This established the high quality demixing was nontrivial. Taken together, these data demonstrate that Bedscales can monitor respiratory status of multiple individuals sharing a bed.

### Ballistocardiographic monitoring

The mechanical force of each heartbeat reverberates through the aorta leading to a characteristic signal known as the ballistocardiogram (BCG) with characteristic waves and segments representing the mechanical correlate of the electrocardiogram (*26*). Close examination of the raw sensor data revealed the BCG was superimposed on the respiratory signals (**Fig. 3A**). Analysis of the frequency spectrum confirmed that energy was concentrated in respiratory and BCG bands (**Fig. 3B**). Therefore, using the sensor with the highest BCG energy, we performed frequency-dependent filtering at cutoffs of 1 Hz and 50 Hz, moving variance and moving mean filtering, and a final frequency-dependent filter with cutoffs of 1 Hz and 3 Hz to create a single-peaked BCG-derived signal that could span both normal and elevated heart rates (**Fig. 3C**). This allowed longitudinal adherence-independent monitoring of cardiac rate and regularity and allowed inference of relative cardiac ejection forces. We validated the Bedscale heart rate estimations by comparing to simultaneously recorded electrocardiograms over 15 minutes. Bland-Altman plots revealed close quantitative agreement between the Bedscale and ECG (SD = 0.9 bpm) (**Fig. 3E, fig. S7**). Quantification of inter-beat interval variability allowed discrimination of patients with atrial fibrillation (SD = 0.17 seconds) and normal sinus rhythm (SD = 0.11 seconds) (**Fig. 3F**). Since respirations are known to alter stroke volume via changes in intrathoracic pressure, we examined the magnitude of the single-peak BCG signal as a function of respiratory phase. Indeed, the ratio of the inspiratory to expiratory single-peak BCG amplitude was systematically greater than 1, indicating respirophasic stroke volume variation resulting from cardiopulmonary coupling (**Fig. 3G**) (*27, 28*).

**Figure 3.**
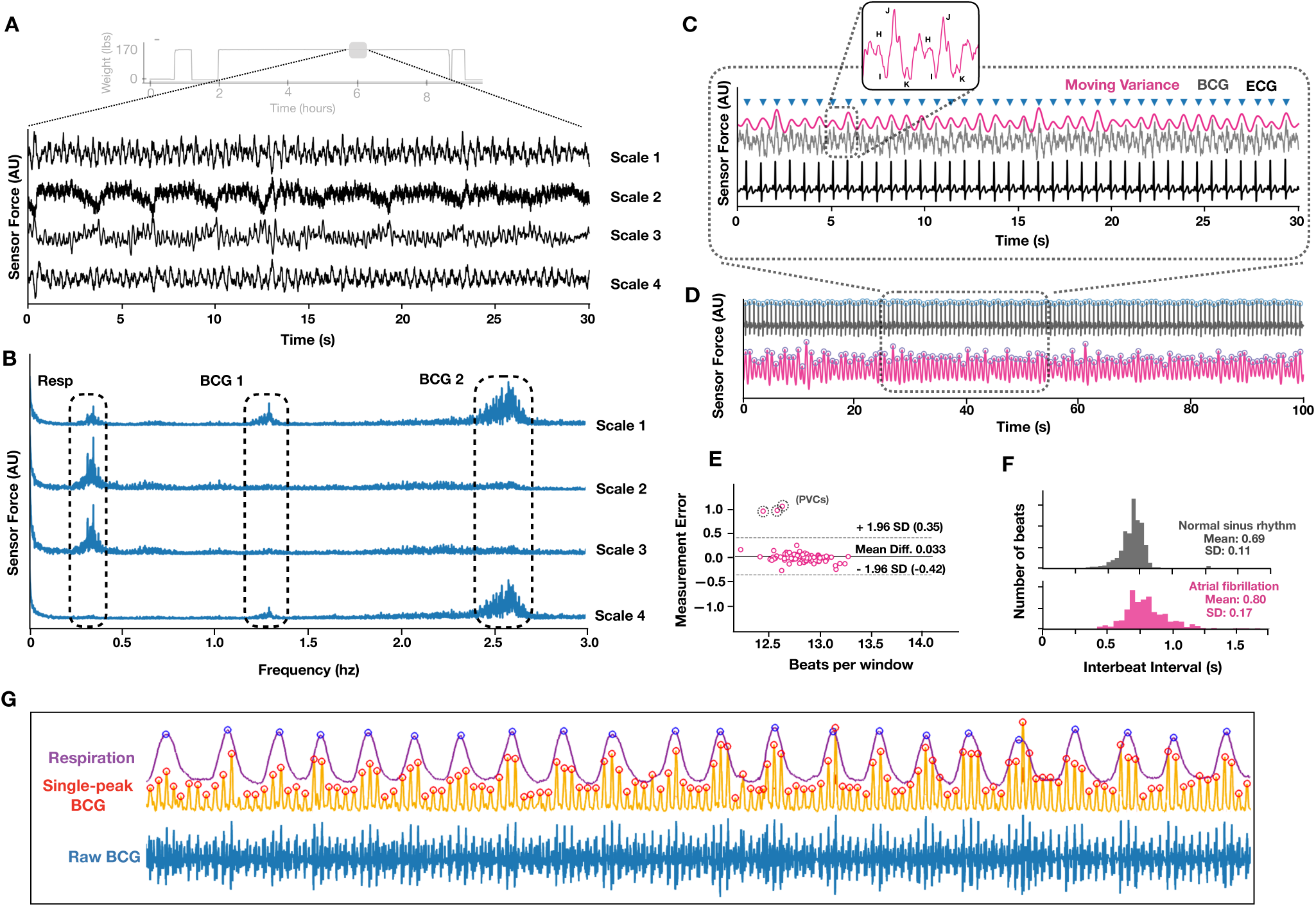
Adherence-independent longitudinal ballistocardiographic monitoring using Bedscales. (**a**) Raw signals (black) from each of the 4 legs during an overnight recording (light grey is the overnight weight signal). (**b**) Frequency spectrum from each of the 4 legs highlighting respiratory and BCG frequency bands and showing the distribution of energy across the 4 sensors. (**c**) Single-peak BCG signal (pink) with annotation of peaks (blue) derived from frequency-dependent, moving variance, and moving mean filtering of the underlying BCG (grey). Inset shows annotation of the characteristic BCG peaks. Corresponding ground truth ECG is shown in black. (**d**) Comparison of ECG (black) and single-peak BCG (pink) during correlation study. (**e**) Bland-Altman plot comparing ECG and BCG moving average heart rate (window size 10 seconds, shift 10 seconds). (**f**) Comparison of variability of interbeat intervals from a patient with atrial fibrillation (bottom, SD = 0.17 seconds) compared to one with normal sinus rhythm (top, SD = 0.11 seconds). (**g**) Relationship between respiratory signal (purple with blue annotated peaks), and single-peak BCG (yellow with orange annotated peaks), with underlying BCG (blue) illustrating respirophasic variation is BCG amplitude.

### Sleep study and apnea monitoring

We installed the Bedscales beneath the legs of a hospital bed during overnight sleep studies, which provided an opportunity to measure pathologic respirations from an individual who had a high burden of central and obstructive sleep apneas (OSA and CSA) (**Fig. 4A**). OSA is characterized by anatomical airway obstruction despite ongoing respiratory effort, whereas CSA is characterized by repetitive cessation of respiratory air flow during sleep due to lack of ventilatory effort; both are common in patients with HF (*29-31*). During the ∼8-hour study, we longitudinally measured respiratory signals from the Bedscales along with the more cumbersome chest and abdomen belts and nasal pressure transducers used as airflow monitors. We aligned and registered data to adjust for differences in sampling frequencies and then quantified the distribution of interbreath intervals greater than 10 seconds and defined these as apneas (**Fig. 4B**). We confirmed that no flow was detected during 407 apnea episodes with mean duration of 22 seconds, a standard deviation of 10.5 seconds, and a maximum apnea duration of 81 seconds. The distribution of apneas was periodic with 5 apnea-dense clusters separated by apnea-free intervals (**Fig. 4C**). Within each apnea cluster we observed substructure during which the longest apneas were followed by the longest apnea-free periods (**Fig. 4D**). Close examination of the tracings demonstrated the Bedscales could discriminate obstructive and central apneas based on the presence of low amplitude unproductive respiratory efforts (obstructive) or the absence of effort (central) (**Fig. 4E-F**). Examination of simultaneous BCGs showed stable amplitude signals in the absence of respiratory effort followed by transient increases in BCG amplitude following the strong negative intrathoracic pressure, providing a new tool with which to investigate beat-by-beat hemodynamic consequences of central and obstructive apneas (**Fig. 4G-H**) (*32*). Taken together, these data demonstrate that Bedscales can perform high fidelity monitoring of normal and pathologic respiratory dynamics and their hemodynamic consequences without the need for obtrusive adherence-dependent sensors and may allow longitudinal characterization of recently reported night-to-night variability (*33, 34*).

**Figure 4.**
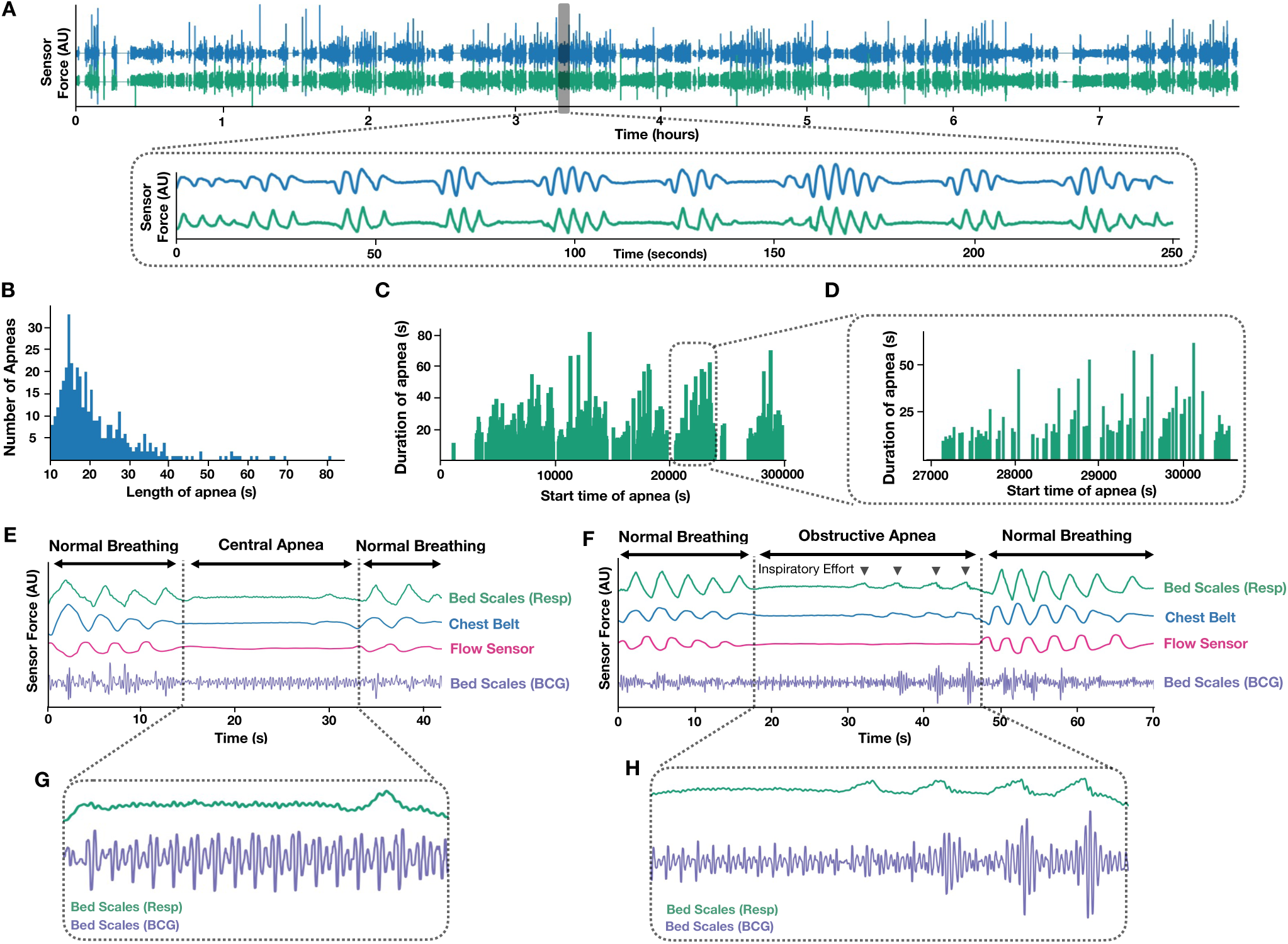
Bedscale measurements during a sleep study in a patient with mixed obstructive and central sleep apnea. (**a**) Bedscale non-contact respiratory signal (green) compared to chest respiratory belt during overnight sleep study. Inset shows an example of this patient’s frequent apneas. (**b**) Histogram of all apneas including all interbreath intervals greater than 10 seconds. (**c**) Duration of apneas vs timing of apneas throughout the overnight study. (**d**) High temporal resolution from one of the five apnea clusters during the night. (**e**) Central sleep apnea episode and (**f**) obstructive sleep apnea episode comparing Bedscale respiratory signal (green) with chest belt (blue), nasal flow sensor (pink) and Bedscales BCG (purple). (**g-h**) Insets show detailed Bedscale respiratory and BCG signals during a (**g**) central apnea (no respiratory effort) and (**h**) an obstructive apnea (low amplitude respiratory effort against a closed airway).

### Long-term in-home monitoring of a patient with heart failure

To evaluate real-world utility of the Bedscales, we performed 3 months of in-home continuous monitoring of a 61-year-old male patient with newly discovered heart failure (EF 15%) who was found to have multivessel coronary artery disease requiring coronary artery bypass surgery (CABG). After being discharged from the hospital, Bedscales were installed under his home recliner where he slept each night. The percent time spent in the recliner each day, which ranged 40-60% before surgery, was 0% during his surgical hospitalization, and was significantly increased to ∼80% for a month before gradually declining to his baseline around the same time he began attending cardiac rehab (**Fig. 5A-B**). Although the patient’s weight fluctuated during the 3 months, it did not show large excursions and he was felt to be euvolemic at clinic visits during the 3 months (**Fig. 5C-D**). His respiratory rate trends were locally stable but exhibited gradual changes over weeks (**Fig. 5E**). Compared to “pre-surgery” respiratory rates of 16-18 bpm, his “post-surgery” respiratory rates were significantly elevated, with an average of 25 bpm punctuated by frequent episodes of more extreme tachypnea (30-40 bpm) (**Fig. 5E-G**). His respiratory rate gradually decreased over 2-3 weeks following surgery and stabilized near his baseline respiratory rate of 18 bpm, consistent with previously reported respiratory rate recovery times following cardiac surgery (**Fig. 5E, F, H**) (*35*). Although his ventricular function modestly improved from 15% to 25%, it remained severely depressed. Consistent with his persistent ischemic cardiomyopathy was his high burden of periodic breathing (periodicity of >30 seconds) and low amplitude nadirs along the spectrum of heart-failure-associated Cheyne-Stokes Breathing (**Fig. 5I**). This suggests Bedscales may enable screening for sleep disordered breathing and provide longitudinal data about relationships between changes in nocturnal breathing and cardiovascular function (*36*). Since development of CSA corresponds to paroxysmal nocturnal dyspnea, Bedscales may also improve detection of HF symptoms, as this often goes unrecognized by patients and unreported to physicians. Throughout the in-home studies, patients reported no concerns and reported being unaware of the sensors.

**Figure 5.**
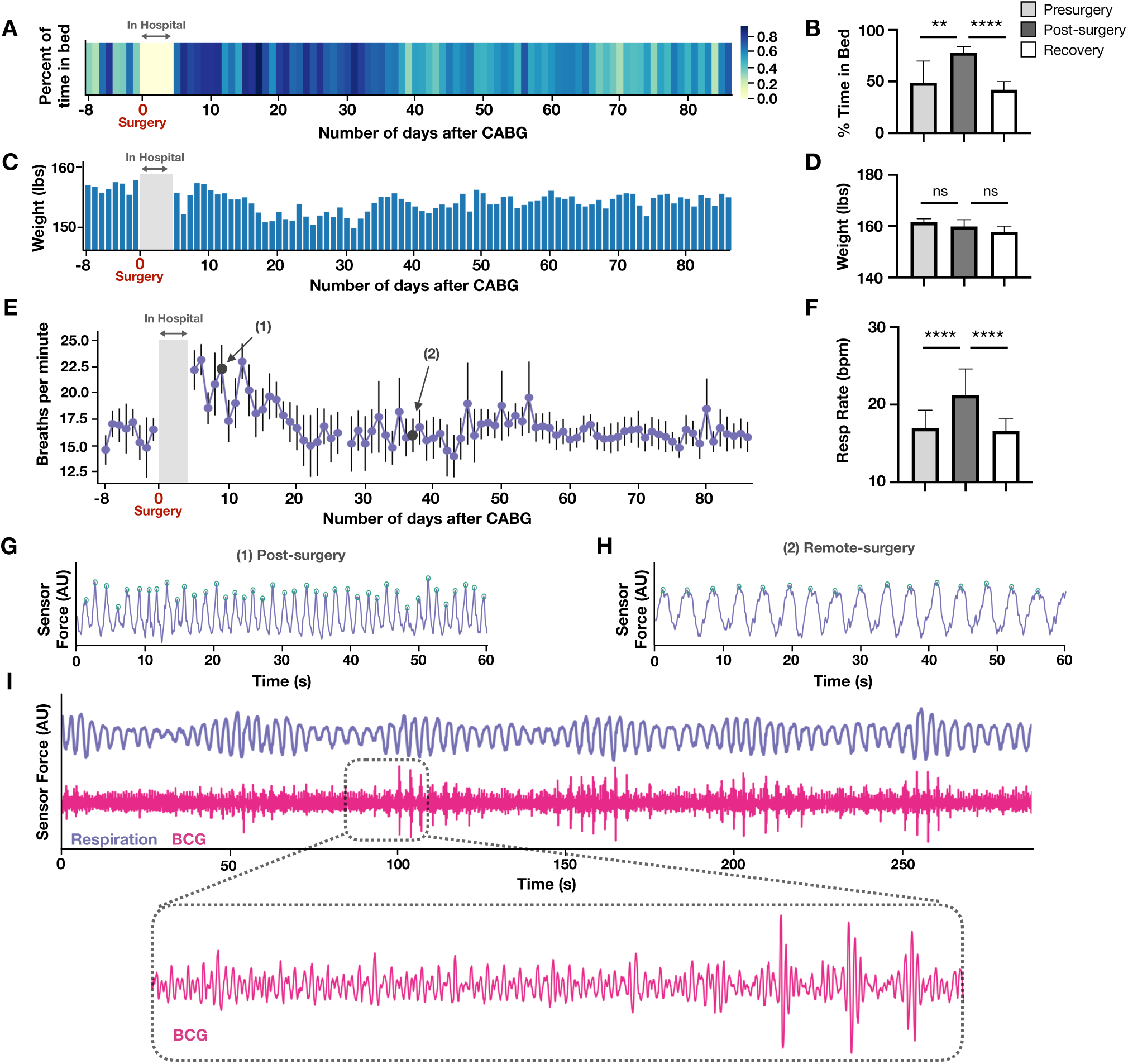
In-home Bedscale measurements in a patient with heart failure before and after cardiac surgery. A 61-year-old male was found to have heart failure and multivessel coronary artery disease requiring bypass surgery. Bedscales were installed under his recliner at home where he sleeps. (**a**) Heatmap shows percent time-in-bed (%TIB) from days 1-7 before surgery, and ∼3 months after surgery. Comparison of pre-post- and remote-surgery %TIB. (**c**) Bar plot of total body weight (TBW) measured by the Bedscales over the course of the 3-month in-home monitoring. (**d**) Comparison of pre-post- and remote-surgery TBW. (**e**) Respiratory rate (RR) average during sleep as measured by the Bedscales before and ∼3 months after surgery. (**f**) Comparison of pre-post- and remote-surgery RR. (**g**) Example of tachypnea (>40 bpm) in the early post-surgical period (black circle annotated as (1)). (**h**) Respiratory rate (16 bpm) one month after surgery (black circle annotated as (2)). (**i**) Periodic respirations suggestive on the spectrum of Cheyne-Stokes breathing and corresponding BCG measured by Bedscales in this heart failure patient with a presurgical ejection fraction of 15% that improved to 25% after surgery. Data in B, D, F, are shown as mean +/- standard deviation. ** *P* <0.01, *** *P* <0.001, \*\*\**P*<0.0001, Mann-Whitney test.

In summary, our results introduce Bedscales, a non-contact adherence-independent home health monitoring system that longitudinally quantifies dynamic forces across diverse amplitudes and time scales to measure weight, respirations, and ballistocardiograms each night, as people sleep in the comfort of their home beds. We solved the “two-body problem” so that weights and respiratory signals can be demixed even when users share the bed with a partner or pet. Finally, we made the technology scalably manufacturable and thus inexpensive compared to implantable medical devices intended for the same purpose (**fig. S8**). This will enable future work focusing on clinical translation and disease-specific applications. In a healthcare environment that is transitioning from fee-for-service to value-based care, Bedscales technology has the potential to make outpatient chronic disease management a data-driven science and in doing so, achieve the triple aim of improving patient satisfaction, improving quality and access for populations, and reducing health care costs (*37*).

## Acknowledgements

We thank Mr. Sean Pawlicki for technical assistance and Ellisa Cholapranee for assistance with sensor testing. The work was funded by NIH-NHLBI T32HL007604, HMS LaDue Fellowship, BU CFTCC Pilot Grant Award (K.R.K.), AHA15MCPRP2569003 (K.R.K), and NIH-NHLBI 1K99HL129168 (K.R.K.).

## Author Contributions

N.H. and K.R.K. designed the device, directed the project, and wrote the initial manuscript. N.H., Z.W., B.H., T.P.C., K.R.K. developed data analysis algorithms and software. P.N.D. and R.L.O. coordinated sleep studies. Q.B. coordinated the in-home heart failure study. N.H., D.L.B., A.S.D., R.L.O, T.P.C. and K.R.K. designed experiments, analyzed data, and edited the manuscript.

## FIGURE CAPTIONS

**Supplementary Figure 1.**
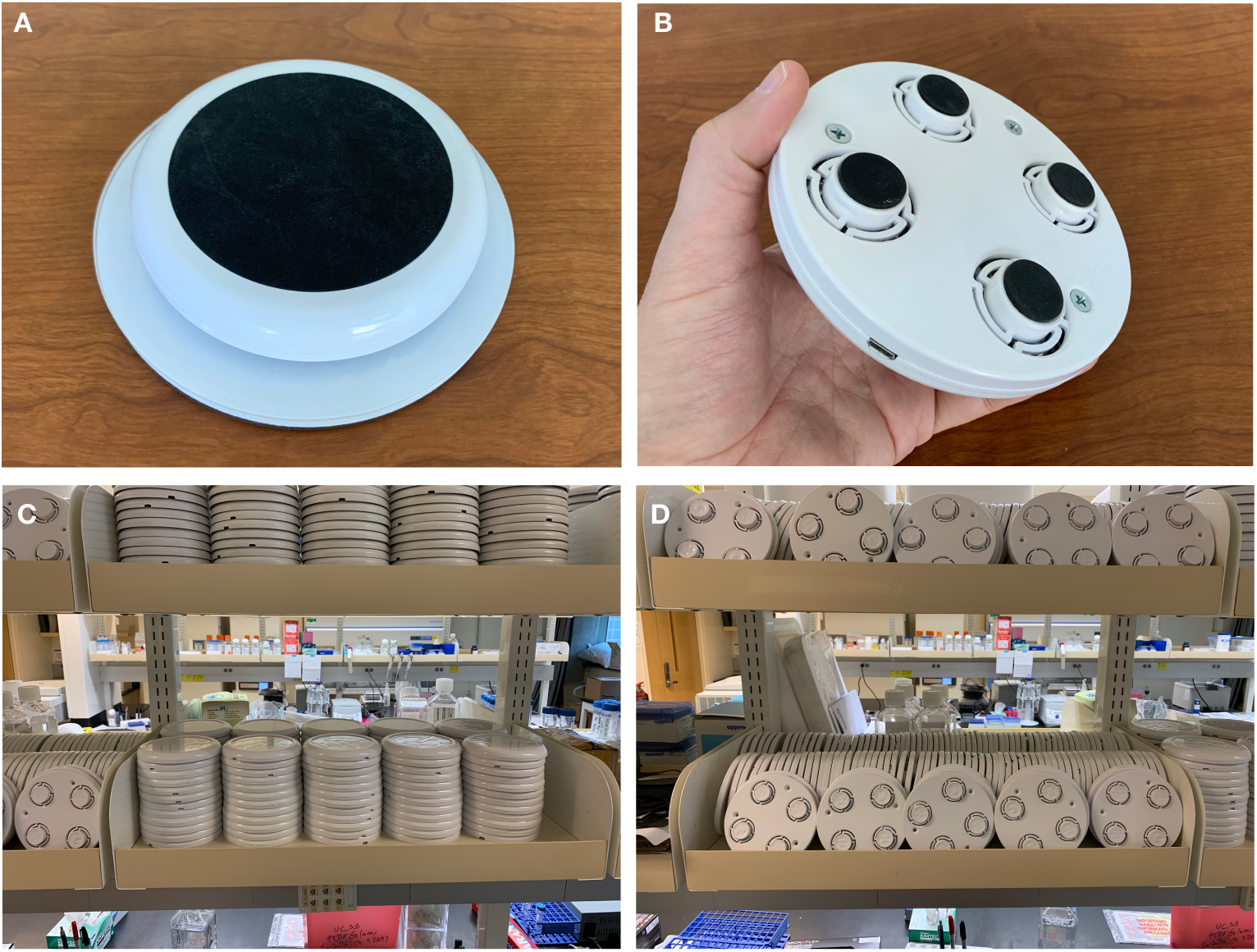
Images of the Bedscale hardware. A fully constructed Bedscale sensor (**a**) top view and (**b**) bottom view. Images of the mass-producible injection molded housing (**c**) plastic tops and (**d**) plastic bottoms.

**Supplementary Figure 2.**
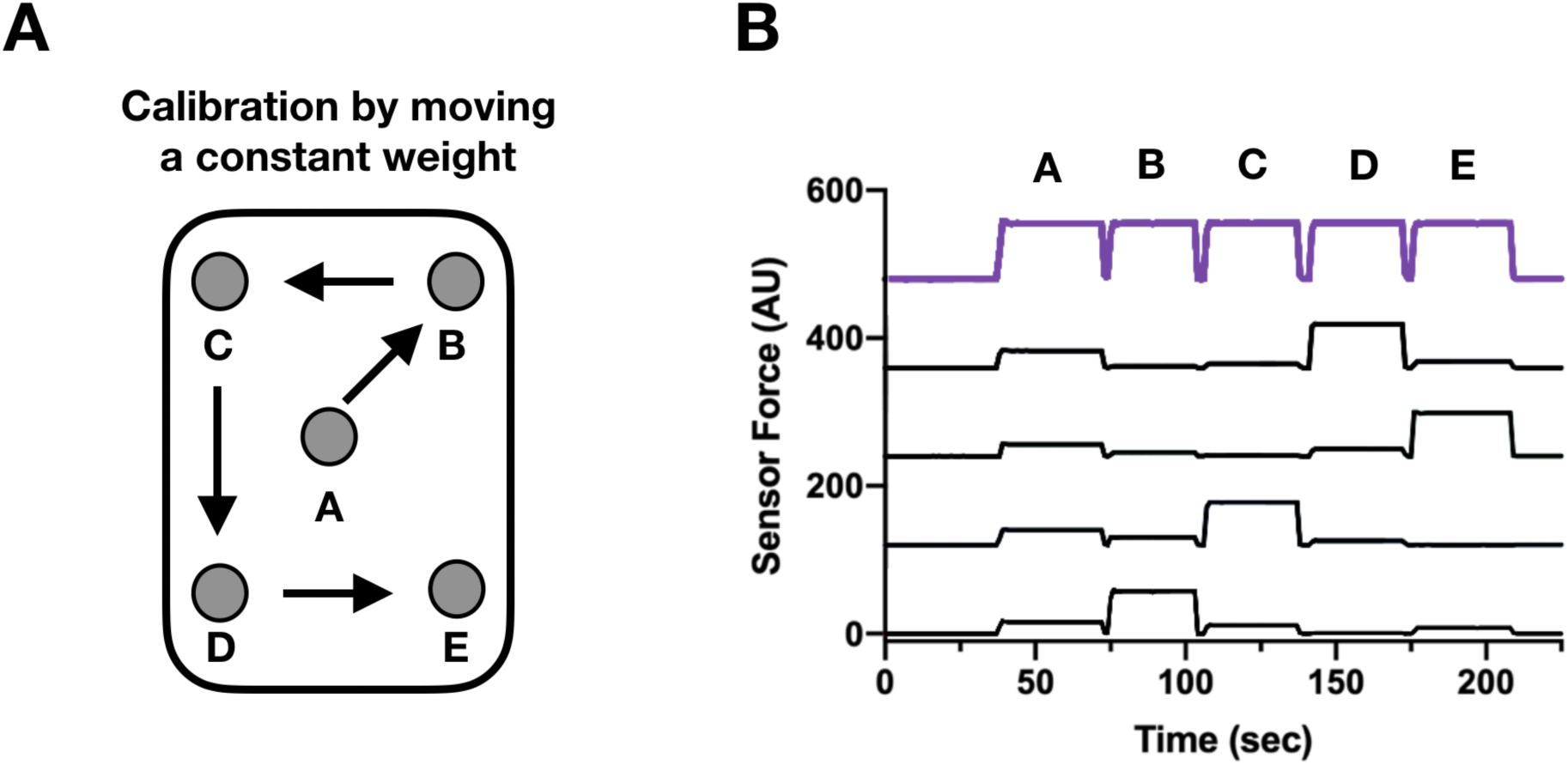
Redistribution of a constant load on a bed. (**a**) Movement of load to multiple positions on a bed (A -> B -> C -> D -> E). (**b**) The black curves show the load measured by each of the four individual sensors beneath each of the four bed legs. The purple curve shows the sum of the loads measured by the four sensors.

**Supplementary Figure 3.**
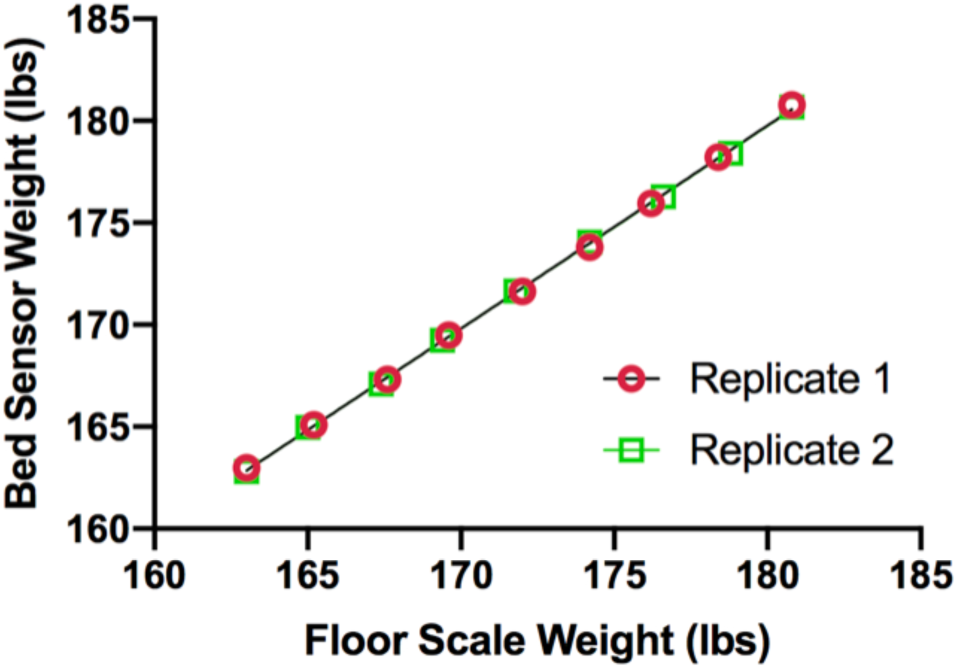
Comparison of Bedscale weights and a commercial floor scale for a person of increasing weight. Plot of floor scale vs Bedscale weight measurements for the individual wearing increasing ankle weights.

**Supplementary Figure 4.**
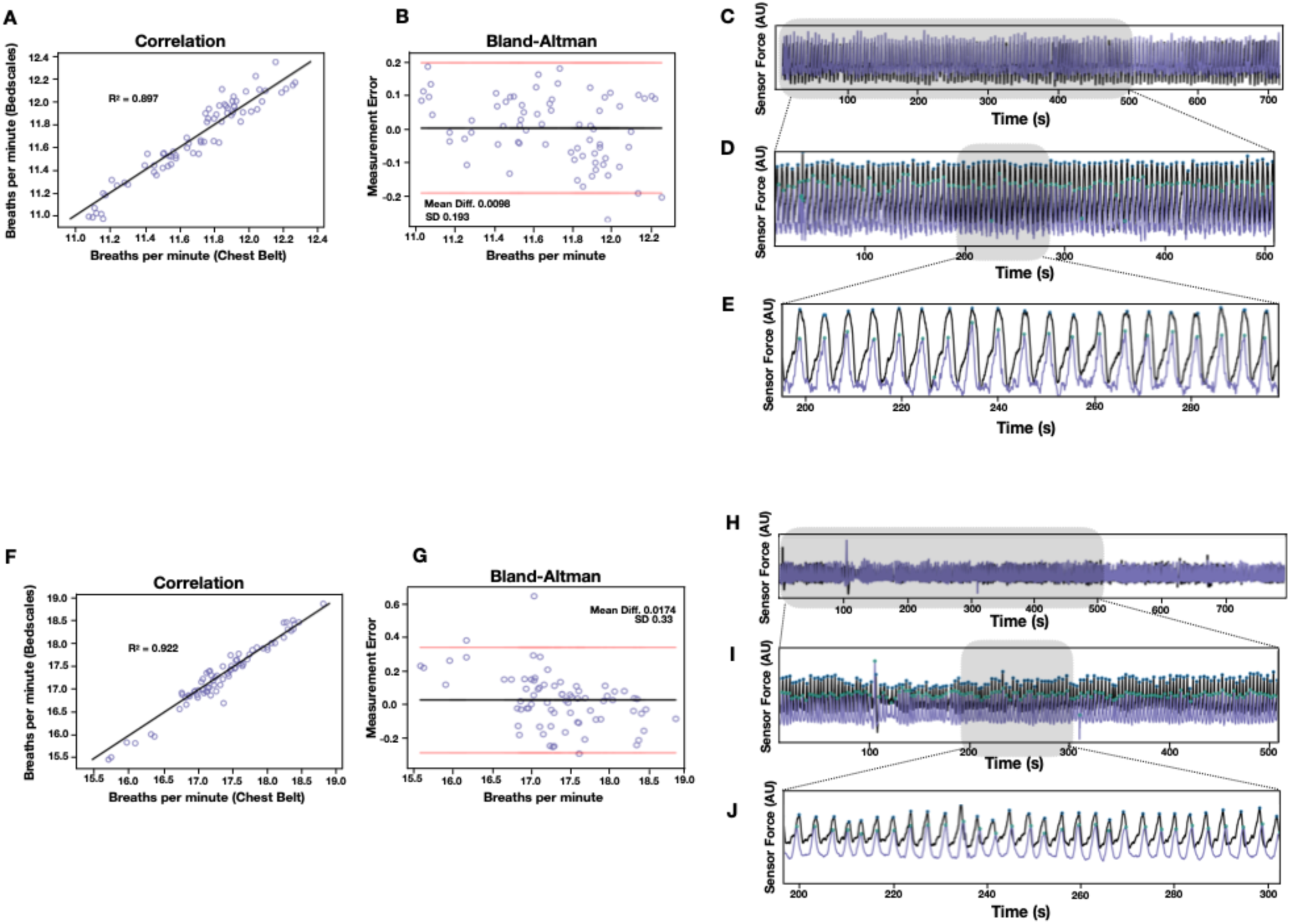
Examples of overnight sleep studies comparing Bedscales and commercial sensors for patients 25 (**a-e**) and 26 (**f-j**). (**a**,**f**) Correlation plots, (**b**,**g**) Bland-Altman plots, (**c-e, h-j**) comparison of respiratory signals from the Bedscales (purple with green annotated peaks) and the commercial chest belt (black with blue annotated peaks) across various time-scales.

**Supplementary Figure 5.**
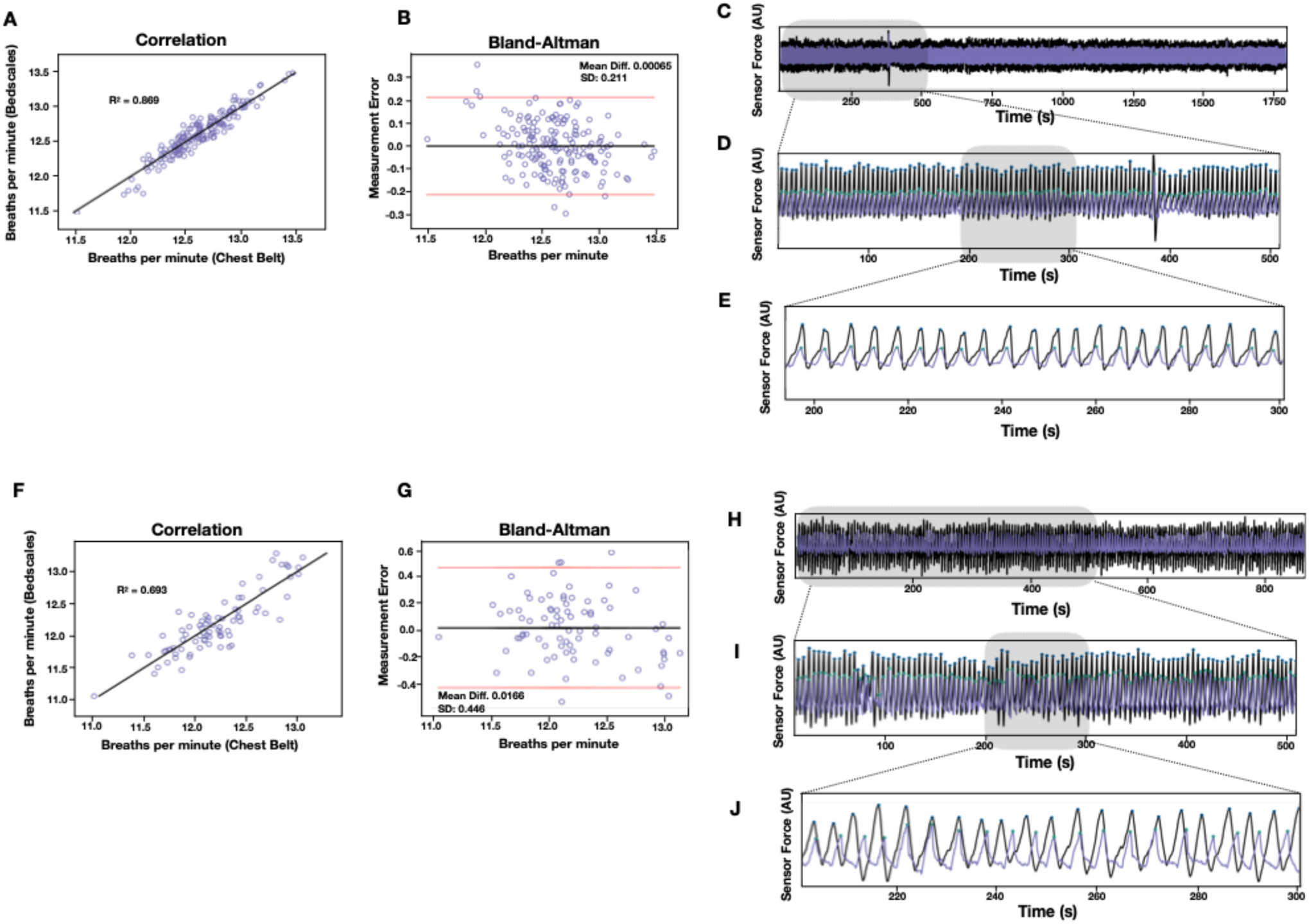
Examples of overnight sleep studies comparing Bedscales and commercial sensors for patients 27 (**a-e**) and 28 (**f-j**). (**a**,**f**) Correlation plots, (**b**,**g**) Bland-Altman plots, (**c-e**,**h-j**) comparison of respiratory signals from the Bedscale (purple with green annotated peaks) and the commercial chest belt (black with blue annotated peaks) across various time-scales.

**Supplementary Figure 6.**
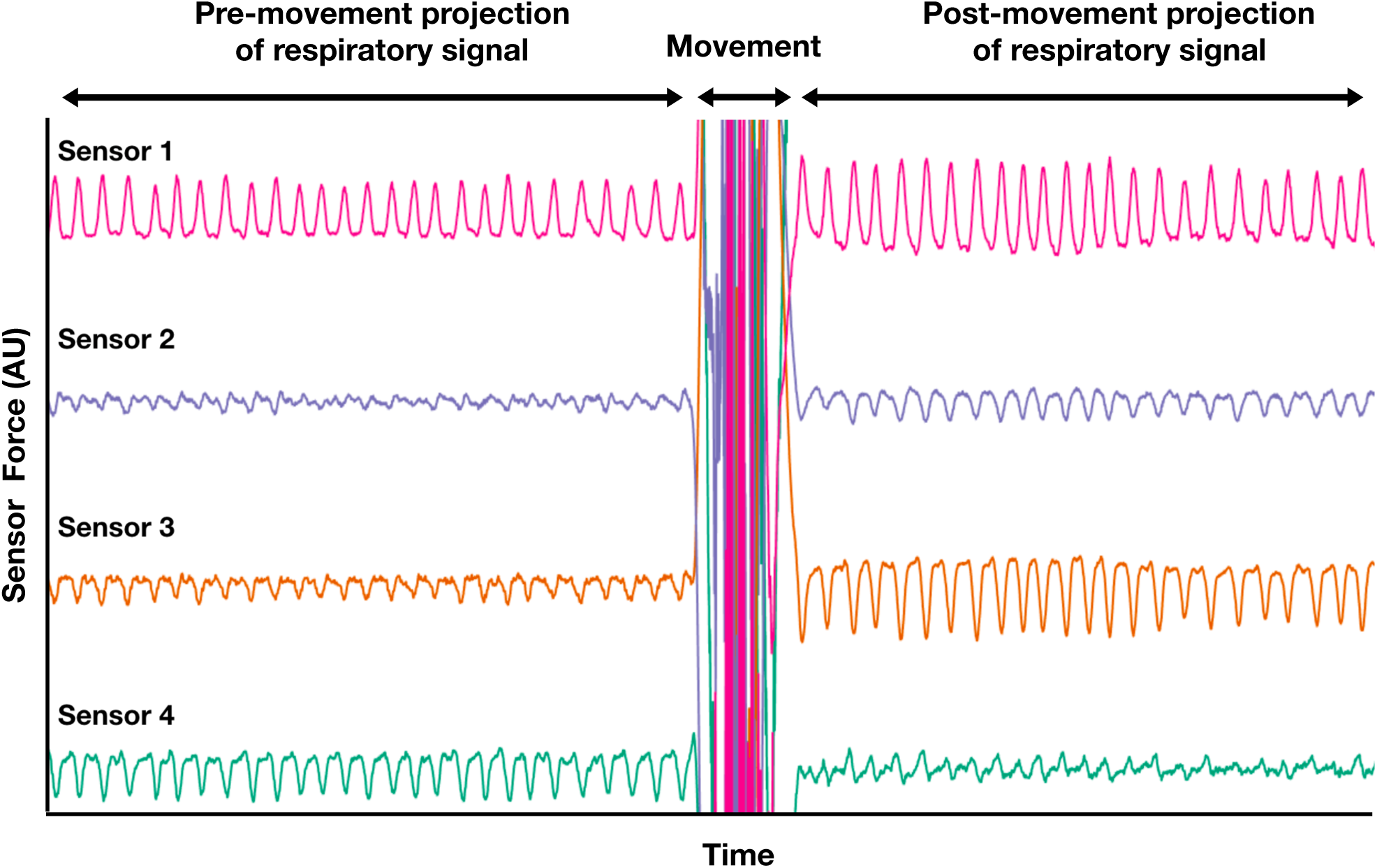
Respiratory signal measured by each of four bed legs during sleep, before and after a movement. The relative amplitudes of respiratory signal measured by the four sensors is locally consistent but changes suddenly after a movement. This indicates that during sleep and between movements, patients can be modeled as respiratory point sources.

**Supplemental Figure 7.**
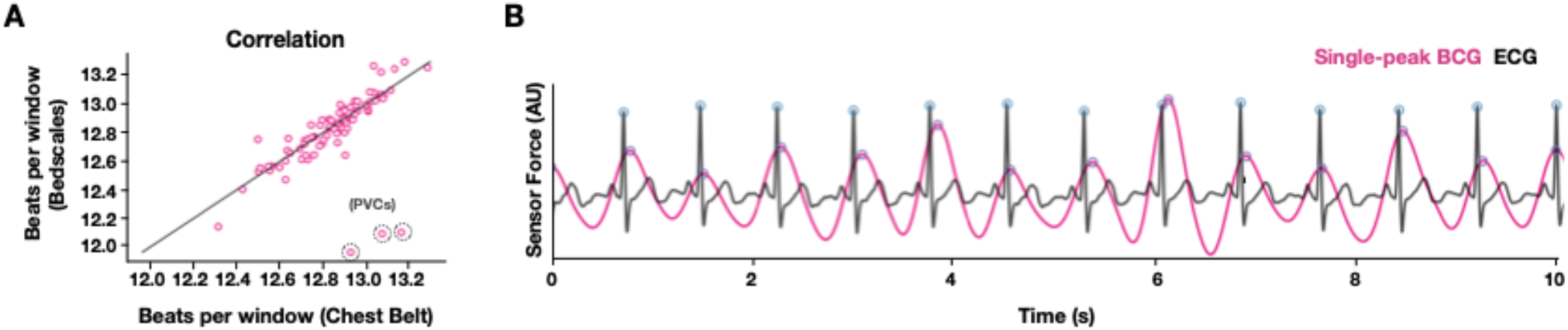
Bedscale BCG-based heart rate estimation. (**a**) Correlation plot with errors due to PVCs circled. (**b**) ECG (black) and corresponding single-peak BCG (pink).

**Supplementary Figure 8.**
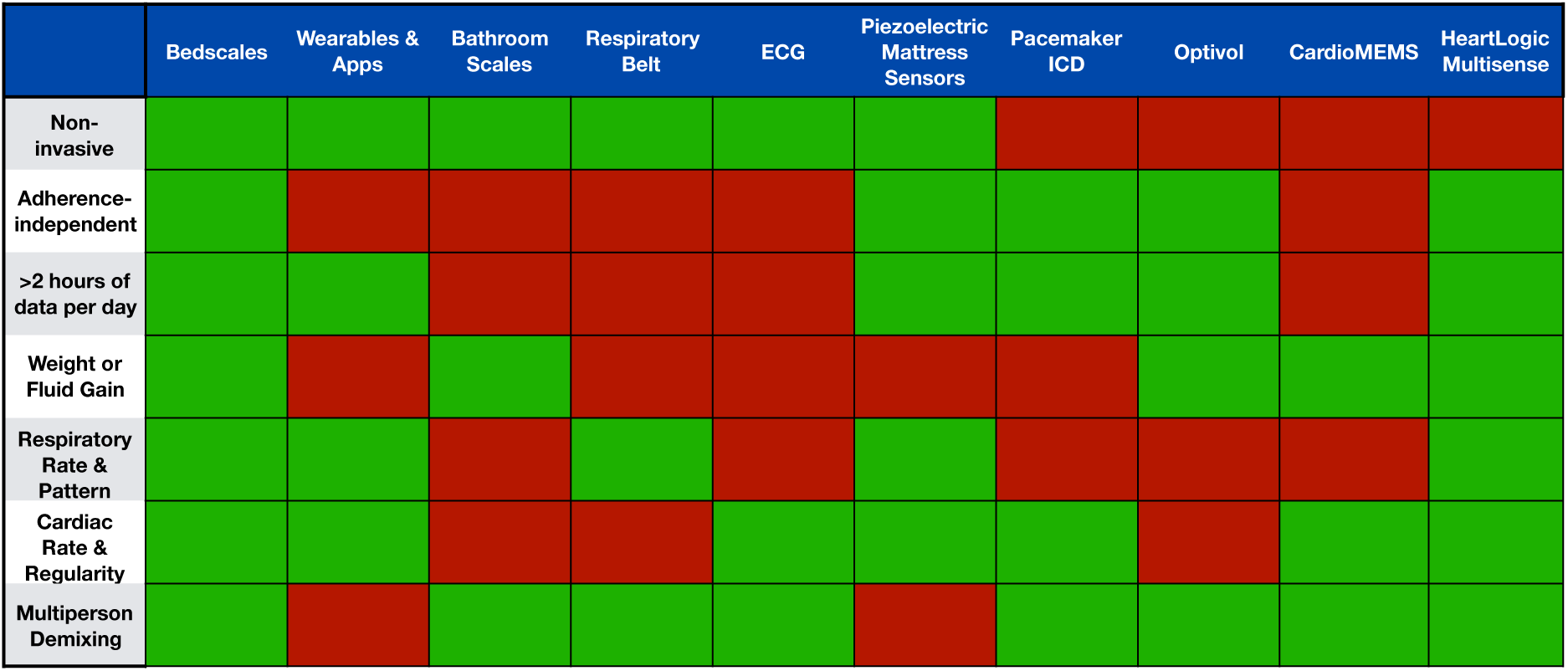
Comparison of features of Bedscales compared to existing heart failure monitoring technologies. For each platform, the features that are available are indicated in green while the features that are unavailable are shown in red.

## MATERIALS AND METHODS

### Bed Sensor Design and Construction

Custom housing was designed using Solidworks (Waltham, MA). Tooling was machined and injection molded parts were manufactured by S. Pawlicki. 50 kg strain gauges were purchased from Sparkfun (Niwot, CO) or a comparable vendor. Custom circuit boards were designed in CircuitMaker (Altium) and featured Hx711 integrated circuit technology (Avia Seminconductor) with a gain of 64 and a sampling frequency of ∼80 Hz. The circuit boards communicated via microUSB to a Raspberry Pi which received electrical power from the building wall outlet, which in turn powered each sensor. The transducers and custom circuit were snap fit into the housing and secured with screws. Rubber feet and tops were die-cut from one-sided adhesive rubber sheets. The Raspberry Pi communicated data using the subject’s home WiFi or a 3G cellular dongle (Hologram, Nova) and transmitted data to Amazon Web Services where it was stored for asynchronous analysis. All analyses were performed in Matlab (Mathworks, Natick, MA) and Python.

### Weight Measurement

Bed sensor weight validation was performed by having a healthy volunteer place increasing numbers of 2.5lbs ankle weights onto their legs and measuring their total body weight on a commercial bathroom scale (Etekcity Digital Body Weight Bathroom Scale) and on the Bedscales bed sensor under a 4-leg twin sized bed. The limits of sensitivity were tested by placing the sensors beneath a 4-leg couch and placing an empty glass on a flat cutting board. At 20 second intervals, 15mL (0.033lbs) aliquots of water were added. The glass was removed and replaced with and without water, and a smartphone was repeatedly added and removed. Weights were calculated by subtracting the total load measured after and before the person got into bed. The total load was estimated by the sum of individual loads measured by the individual sensors beneath each bed leg.

### 2-Person Weight Demixing

The weights of persons sharing a bed were determined by measuring the calibrated sum of all sensors across time and extracting the large differential weight changes that resulted in an alteration of the physiology signal on either side (either transitioning from no respirations to 1 subject or 1 subject to 2 subjects). The weight changes were then classified into two groups termed person 1 and person 2. Simultaneity tests were performed by instructing 2 persons to get into and out of bed at specified temporal intervals in the following sequence - Person 1 (IN), Person 2 (IN), Person 1 (OUT), Person 2 (OUT) and then repeated but exchanging Person 1 and 2. To explore the limits of simultaneity that would still permit decoupling of person weights, we systematically decreased the interval between Person 1 then 2 (or 2 then 1) getting into and out of bed and repeated the experiment for several time intervals (30 sec, 15 sec, 10 sec, and 5 sec), until the maneuver could not reach a steady position in the allotted interval.

### Respiratory Measurement

Bedsensor respiratory signals were created by frequency-dependent filtering with cutoffs of 0.167 Hz and 1.5 Hz. Individual sensor respiratory signals were linearly combined using coefficients derived from the eigenvalues of a moving window PCA analysis. A single respiratory signal was derived by linearly combining the individual sensor respiratory signals weighted by PCA eigenvalues calculated for each 60 second window. Average respiratory rates were calculated by determining the average inter-breath interval during a moving 60 second window using a shift of 30 seconds. Data was aligned and compared to a simultaneously recorded respiratory chest belt quantified using the size moving window.

### 2 Person Respiratory Demixing

Demixing of respiratory signals obtained from 2 simultaneous sleepers was performed using a hidden Markov model. Mechanical respiratory sources were interpreted as latent signals that evolve in a stochastically continuous manner, according to a linear additive Gaussian model, mixed through a linear operation with additive sensor noise to give rise to the signals at the 4 detectors. Interpreting the linear operation as unknown, we used the Expectation-Maximization algorithm to obtain the maximum-likelihood estimate(*38*). Given this estimate, the Kalman smoothing algorithm was used to extract the mechanical respiratory patterns of the two sources(*39*). Validation was performed by simultaneously but independently measuring each respiratory signal using 2 respiratory belts (Vernier, Beaverton, OR). Interbreath intervals were compared by measuring the error between each demixed signal and each ground truth respiratory belt signal. Demixing was deemed successful when the error between each demixed Bedscale signal was high with respect to non-corresponding respiratory belt but low with respect to the corresponding respiratory belt.

### Ballistocardiographic Measurement

We chose the sensor with the greatest energy in the expected frequency bands (BCG1 and BCG 2 in **Fig. 3**). Single-peak BCGs were derived by frequency-dependent filtering (Butterworth, 1^st^ order) with cutoffs of 1 Hz and 50 Hz. This signal was then smoothed using moving variance and moving mean filters and another frequency-dependent filter was applied (Butterworth, 5^th^ order) with cutoffs of 1 Hz and 3 Hz. Average heart rates were calculated by determining the average inter-beat interval during a moving 10 second window using a shift of 10 seconds. Data was compared to a simultaneously recorded ECG. To discriminate atrial fibrillation and normal sinus rhythm, we compared the standard deviations of distributions of BCG-derived interbeat intervals. Respirophasic BCG data was calculated for each peak by measuring the magnitude of the tallest single peak BCG for a beat starting during inspiration divided by the magnitude of the tallest single peak BCG for a beat starting in the expiration phase of the prior breath.

### Clinical Sleep Study

Bedscales were installed beneath the legs of a conventional hospital bed in the Clinical and Translational Research Institute where overnight sleep studies were conducted. The study was performed in accordance with IRB # 171480 and IRB # 180160. As part of another ongoing study, subjects underwent standard in-laboratory polysomnography (PSG) with electroencephalogram (EEG), electro-oculogram, submental and leg electromyogram for sleep staging; nasal pressure and thermistor for airflow measurement; thoracic and abdominal piezoelectric bands for respiratory effort; arterial oxygen saturation monitoring at the finger; and electrocardiogram monitoring for safety. Patients slept supine. Sleep state, arousals, and respiratory events were scored by a registered sleep technologist according to standard American Academy of Sleep Medicine 2012 Recommended Criteria. Signals from the thoracic piezoelectric band and the Bedscales were aligned using custom python scripts. Apneas identified by selecting regions of exceptionally low variance. The timing and duration of each apnea region was recorded.

### In-home Testing

Bedscales were delivered to and installed under the patient’s recliner where he told investigators he slept each night. Signals were recorded locally and transmitted every 24 hours via his home WiFi to a secure Amazon Web Services S3 bucket. Percent time in bed was calculated as the percent of each 24 hour day that the sum of the Bedscales was above a defined threshold. Weights were calculated by creating a histogram of the whole night based on the most prevalent values and the patient’s starting weight. This allowed estimation of the patient’s weight above the baseline weight of the recliner and its nonhuman contents. To avoid spurious measurements, we averaged multiple time points to perform the weight estimation. Weights were then transformed into pounds using calibration factors determined at the time of installation. To calculate respirations, a frequency-dependent filter (Butterworth, 1^st^ order) with cutoffs of 0.167 Hz and 1.5 Hz was used. These signals were then combined via PCA to subsequently perform peak finding. A moving window of size 60 seconds and shift of 30 seconds was used to calculate the respiratory rate of each window in breaths per minute.

### Statistics

Statistical analysis was performed using custom python scripts or GraphPad Prism software. All data are represented as mean values +/- standard deviation unless indicated otherwise. For two-group comparisons, a 2-tailed nonparametric Mann-Whitney test was used unless otherwise specified. All analysis were unpaired. *P* values are indicated by *P* values less than 0.05 were considered significant and are indicated by asterisks as follows: *p<0.05, **p<0.01, ***p<0.001, ****p<0.0001

